# Efficacy of double-stranded RNA for the large-scale control and prevention of sacbrood virus in *Apis cerana* (Hymenoptera: Apidae) apiaries

**DOI:** 10.1101/2021.05.07.443076

**Authors:** Mi-Sun Yoo, A-Tai Truong, Hana Jeong, Do Hyun Hahn, Ju Seong Lee, Soon-Seek Yoon, Yun Sang Cho

**Affiliations:** Parasitic and Honeybee Disease Laboratory, Bacterial and Parasitic Disease Division, Department of Animal and Plant Health Research, Animal and Plant Quarantine Agency, Gimcheon, Gyeongsangbuk-do, Republic of Korea; Faculty of Biotechnology, Thai Nguyen University of Sciences, Thai Nguyen, Vietnam

**Keywords:** Sacbrood virus, *VP1*, RNA interference, Double-stranded RNA, Honeybee, *Apis cerana*

## Abstract

Sacbrood virus (SBV) infection in *Apis cerana* has caused tremendous damage in India, Thailand, Vietnam, and China since the 1970s. The disease caused by this virus results in colony collapse disorder in *A. cerana* and is also a devastating disease affecting *A. cerana* in South Korean apiaries. It has almost resulted in the elimination of the species. Therefore, control measures for this emerging threat are urgently needed. SBV RNA interference (RNAi) targeting *VP1* was prepared to test the safety and efficacy of protection and treatment in artificially infected larvae and in infected colonies in South Korean apiaries. The efficacy of *VP1* double-stranded RNA (dsRNA) was confirmed for the protection and treatment of infected larvae by increasing the survival rate in comparison with that in untreated larvae. Furthermore, an optimal application procedure was established for the large-scale RNAi treatment of SBV in apiaries. The protection of healthy colonies from SBV by RNAi was demonstrated in 100% of apiaries, and the treatment results showed that after five administrations, the SBV in infected colonies was mitigated to a safe level at which no symptoms of the disease were observed. Importantly, the low cost of dsRNA production in this study enables its application as a specific drug in large scale in South Korean apiculture.

## Introduction

Sacbrood virus (SBV) is a positive-sense and single-stranded RNA virus [1] that was first discovered in *Apis mellifera*, in which it causes little damage [2–4]. However, since propagated to *Apis cerana*, the virus has becoming a devastating pathogen and the main cause of the colony collapse of this honeybee species in Vietnam, China, and the Republic of Korea [5–9]. SBV has now become a prevalent pathogen of honeybees and adversely influences the beekeeping industry.

Although various methods for SBV treatment have been tested, an efficient method that is widely application has not been discovered due to the downsides of the methods used thus far. For example, the herbal extract thymol was used to inactivate SBV in *A. cerana*. However, the side effect of this chemical was a decrease in the survival of larvae reared *in vitro* [10]. The use of silver ion has been attempted in South Korea for the treatment and prevention of SBV in *A. cerana* [5]; however, it was not commercialized because of food safety concerns. In addition, lactic acid bacteria have been observed to immunity enhancing effects. However, their application triggers hygienic behavior among nurse bees to eliminate healthy larvae that receive the yogurt [11].

The capacity of insect cells to take up the environmental and artificial double-stranded RNA (dsRNA) via scavenger receptor-mediated endocytosis has been demonstrated [12–14]. The presence of dsRNA in the cytoplasm activates the small interfering RNA (siRNA) pathway via which the RNase type III enzyme Dicer-2 cleaves the dsRNA into small fragments of 18–24 nt [15]. The cleavage is then carried out by Argonaute-2, which contains an “RNA induced silencing complex” [13, 16]. The species-specific targeting of RNA interference (RNAi) has become a potential approach for the control of insect pests in crops [17], and the capacity of RNAi to inhibit and control honeybee parasites has been reported, such as the small hive beetle, *Aethina tumida* Murray [18] and mite, *Varroa destructor* [19]. RNAi is also a promising therapy for the prevention of honeybee viral pathogens and could be developed as a drug for field application [7, 20–23].

The administration of RNAi specific to SBV of *A. cerana* was demonstrated to be effective at blocking viral propagation and increasing the survival rate of larvae *in vitro;* consequently, it could be a promising method for controlling SBV in *A. cerana* [7, 24]. However, clinical studies on the capability of RNAi for wide application for SBV treatment and prevention have not been reported. Accordingly, the present study was conducted to analyze the efficacy of RNAi for the prevention and treatment of South Korean SBV in *A. cerana*, and a procedure for apiary application was developed. Additionally, this commercially cheap RNAi treatment was produced for wide application in South Korean apiculture.

## Materials and methods

### Isolation of sacbrood virus

Larval samples were pulverized using a mortar and pestle, and the presence of SBV and the South Korean genotype of SBV was confirmed by PCR (Table 1). Then, the samples were filtered using 0.45 μm and 0.2 μm syringe filters. After sonicating for 30 s (2 s on and 3 s off), the SBV samples were purified via sucrose density gradient centrifugation (10–50% range) at 32,500 rpm for 4 h with an SW 41 Ti Swinging-Bucket Rotor (Beckman Coulter Inc., Brea, CA). The SBV band that corresponded to the 40% sucrose gradient was harvested and then centrifuged at 25,000 rpm for 12 h; then, the SBV was eluted with ddH_2_O. Photographs were taken with a transmission electron microscope (H-7100FA, Hitachi Ltd., Tokyo, Japan) to confirm the viral particles of 27.8 ± 0.4 nm in size, which were similar in shape to other picornaviruses (S1 Fig).

**Table 1.**
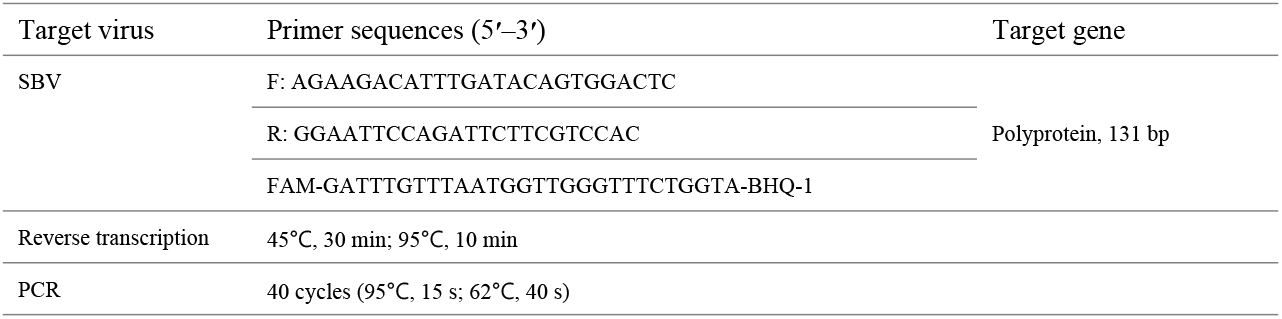
Primers, probes, and PCR conditions for the identification of sacbrood virus (SBV)

### Extraction of viral RNA

The RNA of SBV was extracted from infected larvae using QIAamp Viral RNA Mini Kit (QIAGEN, Hilden, Germany). Five larvae from each colony were collected in 1 mL phosphate buffered saline (PBS) in a tissue grinding tube with 2.381 mm steel beads (SNC, Hanam, South Korea), and 100 μL of the liquid was used for RNA extraction. For RNA extraction from adult bees, two bees of each colony were collected and put in a tissue grinding tube with 2.381 mm steel beads (SNC). After adding 1 mL of PBS solution the bees were homogenized with a Precellys 24 Tissue Homogeniser (Bertin Instruments, Montigny-le-Bretonneux, France), and 200 μL homogenate was used for RNA extraction. The procedure was carried out according to the manufacturer’s instructions. Finally, 50 μL RNA solution was acquired from each sample.

### Construction of SBV recombinant plasmid

The synthesis of SBV VP1 cDNA fragments was carried out using the SuperScript III First-Strand Synthesis System for RT-PCR (Thermo Fisher Scientific, Waltham, MA). A reaction mix (10 μL) consisting of 4 μL of RNA, 5 μL of reverse primer (10 pmol/μL; Table 2), and 1 μL of 10 mM dNTP mix was incubated at 65°C for 5 min and then placed on ice for 1 min. Then, a solution containing 2 μL of 10× RT buffer, 4 μL of 25 mM MgCl_2_, 1 μL of RNaseOUT, 1 μL of SuperScript III RT, and 2 μL of ddH_2_O was added to obtain 20 μL of reaction mixture. Reverse transcription was conducted by incubating the reaction mix at 50°C for 50 min, then at 85°C for 5 min, and, finally, after chilling on ice, 1 μL RNase H was added and the mix was incubated at 37 °C for 20 min. The cDNA was stored at −20°C until further use.

**Table 2.**
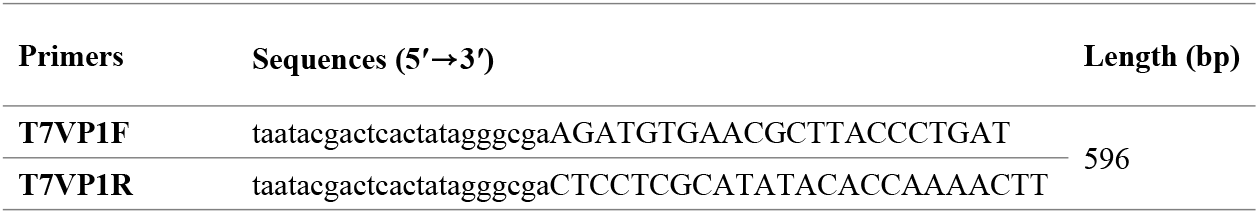
Primers with T7 promoter used for manufacturing double-stranded RNA (dsRNA)

PCR was performed with the synthesized cDNA, the primer set T7VP1F/R for VP1 fragments (Table 2), and TOPsimple DryMIX-HOT PCR premix (Enzynomics, Daejeon, South Korea). The 20 μL PCR mixture was composed of 1 μL of cDNA, 1 μL (10 pmol) of each primer, and 17 μL of ddH_2_O. PCR conditions were as follows: 94°C for 5 min, followed by 40 cycles at 94°C for 15 s, 62°C for 15 s, 72°C for 15 s, and the final extension at 72°C for 10 min. After confirming the presence of the *VP1* band (596 bp; S2 Fig), the DNA fragment was extracted and purified using the QIAquick Gel Extraction Kit (QIAGEN), and a recombinant plasmid carrying the *VP1* fragment was constructed with TOPcloner TA Kit (Enzynomics). The recombinant plasmid was sequenced by GenoTech, Daejeon, South Korea, and compared to the SBV sequences reported in NCBI using the Basic Local Alignment Search Tool (BLAST). There was 99% similarity with the South Korean SBV (NCBI accession no.: HQ322114), and the recombinant plasmid was used as a DNA template for the PCR to produce dsRNA.

### Production of anti-SBV dsRNA

The 596 bp dsRNA corresponding to *VP1* of the South Korean SBV (NCBI accession no.: HQ322114; 1596-2191) was synthesized by transcribing the T7 promoter, which was located on the downstream sequence of primers (Table 2), *VP1* dsDNA produced by PCR served as the template, and the *in vitro* Transcription T7 kit (for siRNA synthesis; No. 6140, TaKaRa Bio Inc., Kusatsu, Japan) was used. The dsRNA synthesis was performed according to the manufacturer’s instructions. Briefly, the reaction mixture was prepared by mixing 10 μL of NTP, 3 μL of 10× reaction buffer, 1 μg of linear DNA template, and 2 μL of T7 Enzyme Mix (TaKaRa). This was adjusted to 30 μL with nuclease-free ddH_2_O before incubating the mixture at 42°C for 2 h. Subsequently, 30 μL of LiCl solution was added for 30 min at −20°C to precipitate RNA, the mixture was ultracentrifuged at 16,000 × *g* for 15 min, and the supernatant was discarded. Thereafter, 1 mL of 70% ethanol was added, and the mixture was centrifuged again at 10,000 × *g* for 5 min. After discarding the supernatant, the RNA was dissolved in 30 μL of elution buffer and then stored at −70°C until further use. A large quantity of dsRNA was produced in collaboration with Genolution, Inc. (Seoul, South Korea) to extend the clinical application to apiaries in the field.

### Real-time reverse transcription PCR for evaluation of SBV inhibition by dsRNA

Real-time reverse transcription PCR (qRT-PCR) was used to quantitatively evaluate the efficacy of dsRNA for inhibiting SBV in infected honeybees. A specific primer pair, namely, KSBV-PCR-F (5’-GACCAAGAAGGGAATCAG-3’) and KSBV-PCR-123R (5’-CATCTTCTTTAGCACCAGTATCCA-3’), was designed to amplify a 123 bp fragment of South Korean SBV (NCBI accession no.: HQ322114; 2306-2428). The PCR was performed with the One Step TB Green PrimeScript RT-PCR Kit II (TaKaRa), and SYBR Green was used as a fluorescent dye for detection. The corresponding 20 μL reaction mix was composed of 2 μL of sample RNA, 1 μL (10 pmol) of each primer, 1 μL of PrimeScript One Step Enzyme Mix (TaKaRa), 10 μL of 2× One Step RT-PCR buffer (TaKaRa), and 5 μL ddH_2_O. The CFX96 Touch Real-time PCR Detection System (Bio-Rad, Hercules, CA) was used for qRT-PCR. Reverse transcription was performed at 42°C for 30 min, followed by PCR at 95°C for 5 min, then 40 cycles at 95°C for 30 s, 55°C for 30 s, and 72°C for 30 s. To identify positive results, the peak melting curves were analyzed. To estimate the number of DNA copies, standard curves representing the relationship between the cycle threshold of amplification and the initial number of DNA copies, namely, from 10^10^ to 10^1^ (10-fold dilution), were established in triplicated PCRs (S3 Fig).

### Efficacy of dsRNA in vitro

The safety and efficacy of dsRNA were evaluated *in vitro* with artificially reared larvae. Honeybee larvae were transferred into the wells of 24-well plates for cell culture, and then incubated at 35°C and 80% humidity. An artificial nutrient solution composed of 6% D-glucose, 6% D-fructose, 1% yeast extract, 33% Gibco Grace’s insect medium (Thermo Fisher Scientific), and 50% royal jelly was provided to each larva at a dose of 100 μL/day.

To assess the efficacy of dsRNA in the inhibition of SBV *in vitro*, we examined the effects of treatment with dsRNA on SBV infection. Three-day-old larvae were artificially infected with SBV. Field larvae that were confirmed to have been infected with SBV, based on clinical signs and via RT-PCR (Table 1), were ground until they were liquefied, and then, they were used to prepare feeding solution by adding other nutrients with ratios similar to those described above. The feeding solution with 10^8^ copies of SBV was supplied to each of the larva being reared in one dose. The experiment was designed with four groups of larvae. Group 1 contained larvae in four 24-well plates (*n* = 96) that were orally administered with dsRNA 1 day after SBV infection. The dsRNA solution was prepared by mixing 1 μg of dsRNA with the feeding solution, and it was supplied to each larva in one dose. The SBV-infected larvae in the other four plates of Group 2 were not fed dsRNA and served as the control. In Group 3, the larvae were fed only dsRNA in feeding solution (no SBV infection) to evaluate the safety of administering dsRNA to larvae *in vitro*. In Group 4, only feeding solution (no dsRNA and no SBV) was supplied. The number of living larvae was counted daily until the 8^th^ day of the inoculation period to calculate the survival rate of larvae.

### Safety and efficacy under controlled conditions

Two hives of *A. cerana* were tested in different isolated rooms. The first hive was sprayed with a feeding solution that was contaminated with SBV. The feeding solution was prepared by mixing 990 mL of ddH_2_O, 600 g of sucrose, 170 g of honey, and 6 mL of apple vinegar. The SBV-contaminated solution was prepared by grinding 25 SBV-infected larvae in 5 mL of feeding solution. The number of RNA copies of SBV in the 5 mL spraying solution was estimated by qRT-PCR to be approximately 3.58 × 10^12^ equivalent copies. The other hive served as a control and was treated with the SBV-free feeding solution. Feeding solution was sprayed to the bees every day, and room temperature was retained at 25°C during the experimental period. Larvae and adults were collected at 1, 5, 8, and 12 days after infection (i.e. day 0), and the level of SBV infection was confirmed by Korean SBV (kSBV) real-time PCR. Five larvae or five adult honeybees were randomly collected at each time for the quantitative detection of SBV, and SBV DNA copy/bee was finally calculated. Adults in the control group were collected at the same time, and their level of SBV infection was compared with that of the honeybees in the SBV infection group.

The other hive of *A. cerana* was placed in another isolated room and was sprayed once with 5 mL of a feeding solution that contained 10 mg *VP1* dsRNA. Three days after the administration of dsRNA, larvae were artificially infected with SBV by spraying them with 5 mL of a feeding solution that contained the same quantity of SBV as described above. The changes in the SBV infection level were measured over time (before infection, on the infection date, and 2, 4, 7, and 10 days after infection) by SBV qRT-PCR.

### Efficacy in the field

The prevention efficacy of dsRNA against SBV was first evaluated in 78 colonies across 17 apiaries, of which 18 colonies were first treated with 10 mg of dsRNA/hive by spraying or oral administration. Afterward, dsRNA application was extended to 60 other colonies at 20 mg/hive via oral administration. Presence of SBV in the colonies was confirmed by qRT-PCR, with 7.77 × 10^5^–2.11 × 10^8^ copies of SBV DNA per larva. However, no clinical sign of SBV disease was observed in the colonies. The hives were administered with dsRNA five times, with a 1-week interval. In addition, seven colonies from four apiaries were not supplied with dsRNA, and these were used as the controls. The apiaries were located near the disease-outbreak regions (approximately 6 km).

Finally, dsRNA was applied for the prevention and treatment of SBV in large-scale in 269 colonies belonging to 33 different apiaries in South Korea, with oral administration of 20 mg dsRNA/hive. In the colonies, the queens were not confined during the period of dsRNA application.

### Data analysis

Survival rates in the dsRNA treatment group and the control groups *in vitro* test were analyzed using the Kaplan-Meier method [25, 26]. The comparison was also performed to verify the efficacy of dsRNA in inhibiting SBV in infected colonies in the field by comparing the SBV DNA copies before and after treatment. The Student’s *t*-test was used to test for significant difference in Microsoft Excel 2019 (Microsoft Corp., Redmond, WA). Probabilities below 5% were considered to indicate significance.

## Results

### Efficacy of dsRNA in vitro

The efficacy of dsRNA for inhibiting SBV was demonstrated in the survival rate of larvae reared *in vitro*. The survival of SBV infected larvae without dsRNA treatment rapidly decreased from 97.5% at the 3^rd^ day of inoculation to 24.0% at the 8^th^ day of inoculation. Meanwhile, the infected larvae that received the dsRNA treatment showed a higher rate of survival (*p* = 0.02), namely, 97.5% 3 and 4 days after inoculation, before decreasing to 87.8% on the 5^th^ day and to 60.5% at the end of the inoculation period (Fig 1). In addition, oral administration of dsRNA was safe for larvae. This was demonstrated by the survival rate of the group that was fed only dsRNA (without SBV) not significantly differing from that of the larvae that only received the feeding solution without SBV (*p* = 0.13; Fig 1).

**Fig 1.**
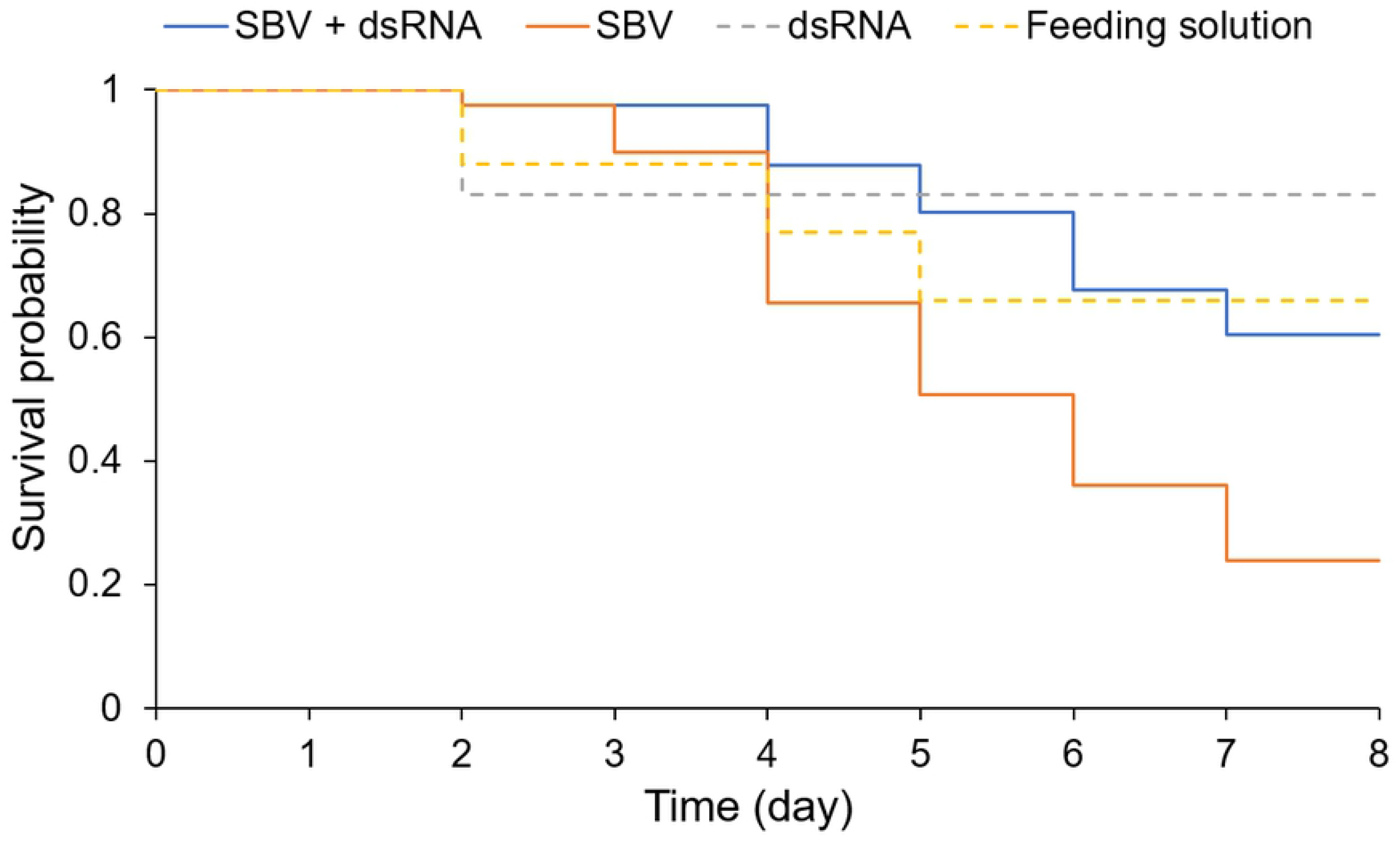
Treatment of sacbrood virus (SBV) using *VP1* double-stranded RNA (dsRNA) in honeybee larvae reared *in vitro*. Survival probability of larvae was calculated according to the Kaplan-Meier method for groups that were artificially infected with SBV and treated with dsRNA (SBV+dsRNA) and not treated with dsRNA (SBV). Two other groups of larvae not infected with SBV were fed dsRNA mixed with feeding solution (dsRNA) and feeding solution only (feeding solution).

### Prevention of SBV disease in controlled conditions

After the artificial infection of *A. cerana* colonies with SBV, the number of SBV DNA copies peaked (approximately 10^10^ copies/bee) 5 days after inoculation and remained at that level until the 12^th^ day after inoculation (Fig 2A). In the colonies that were treated with dsRNA 3 days prior to SBV infection, the development of SBV occurred and remained high (approximately 10^10^ DNA copies/bee) from the 2^nd^ to the 7^th^ day after SBV infection. The capability of dsRNA to inhibit SBV was demonstrated after 7 days of infection by the considerable decrease in SBV to around 10^4^ copies/adult bee and 10^2^ DNA copies/larva by the 10^th^ day following infection (Fig 2B). Afterward, the colony was placed in natural conditions and observed for 6 months, with no SBV disease symptoms observed. In addition, the infected colony with no dsRNA administration showed colony collapse after 2 months.

**Fig 2.**
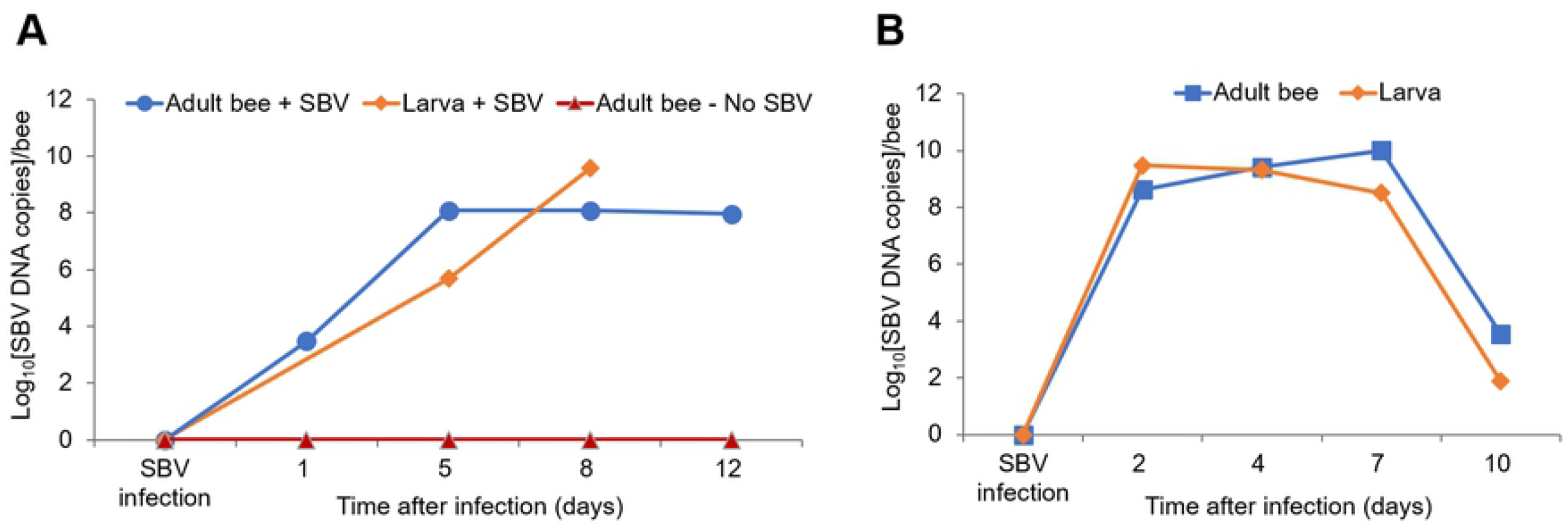
Prevention of sacbrood virus (SBV) using *VP1* double-stranded RNA (dsRNA). The experiment was conducted in two different colonies; qRT-PCR was used to evaluate the efficiency of dsRNA against SBV via the quantitative detection of SBV (i.e. DNA copies). (**A**) Control experiment without dsRNA application was conducted in two colonies: one was artificially infected with SBV, and the increase in SBV in adult honeybees (Adult bee – SBV) and larvae (Larva – SBV) was evaluated, and the other colony was maintained in a healthy condition without SBV infection (Adult bee – no SBV). (**B**) In another colony, dsRNA was supplied to the hive by spraying 3 days before SBV infection, and the SBV in adult honeybees and larvae was quantified 2, 4, 7, and 10 days after infection.

### Field application of dsRNA for the prevention and treatment of SBV

The preventive administration of dsRNA resulted in the absence of clinical symptoms of SBV infection in all 78 *A. cerana* colonies that received dsRNA via spraying or oral administration. The colonies that received doses of 10 mg or 20 mg of dsRNA developed resistance against SBV (Table 3). Meanwhile, clinical symptoms of SBV disease were observed in 3 of the 7 control colonies (42%), which were not administered dsRNA (Fig 3).

**Fig 3.**
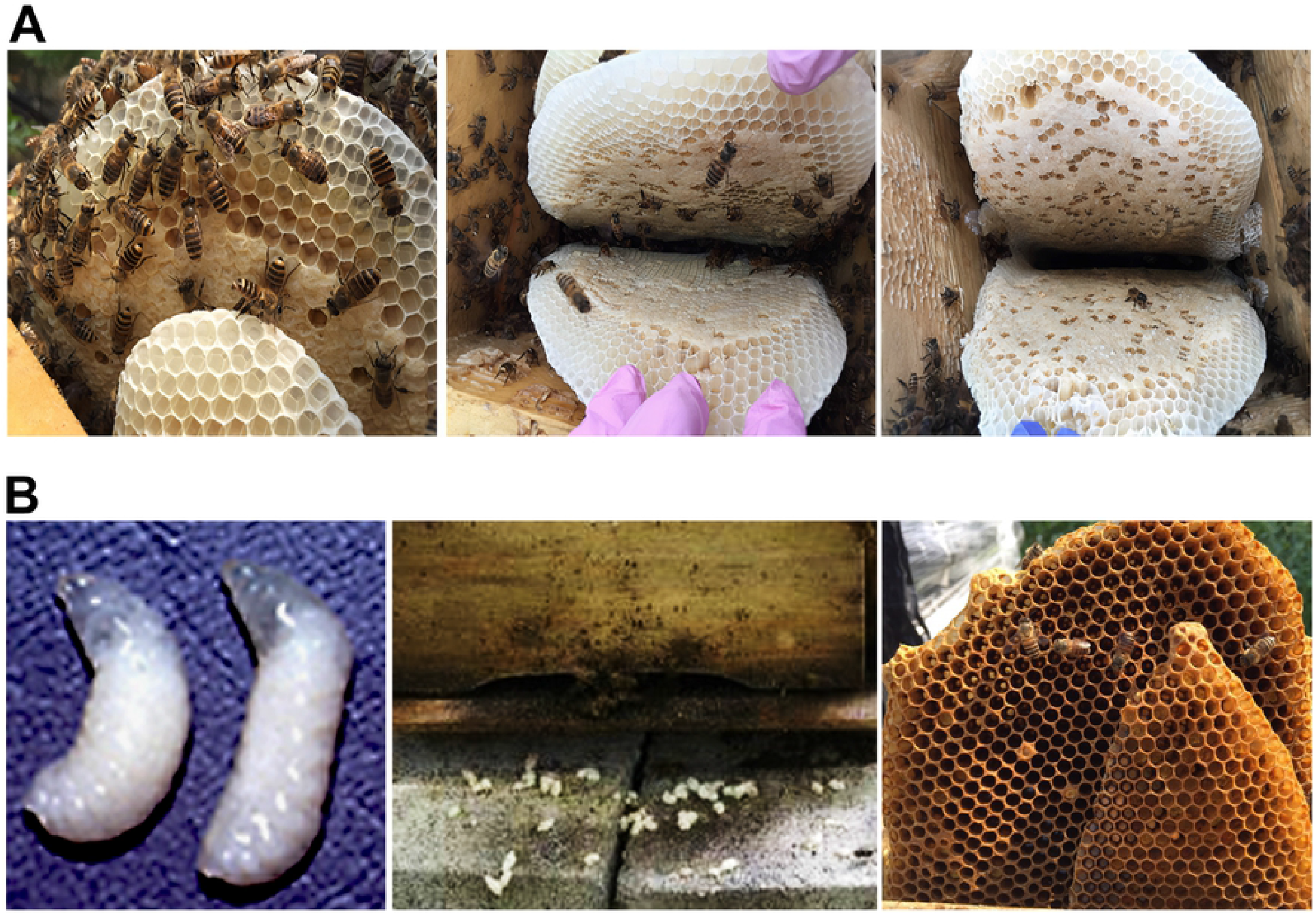
Colonies of *Apis cerana* after sacbrood virus (SBV) infection and double-stranded RNA (dsRNA) treatment. After receiving dsRNA five times (1-week intervals), the SBV-infected colonies showed the healthy condition, and there were no visible symptoms of the disease in the hives (**A**). Meanwhile, the colonies without dsRNA treatment showed clinical signs of SBV disease (**B**).

**Table 3.** Field application of double-stranded RNA (dsRNA) for the prevention of sacbrood virus (SBV)

| Groups | Application route (conc.) | Apiary No. | Colony No. | No. of hives with clinical symptoms |
| --- | --- | --- | --- | --- |
| dsRNA treatment | Spray (10 mg) | 5 | 9 | 0 |
|  | Oral (10 mg) | 5 | 9 | 0 |
|  | Oral (20 mg) | 7 | 60 | 0 |
|  | Subtotal | 17 | 78 | 0 (0%) |
| Negative control | No treatment | 4 | 7 | 3 (42%) |

The quantitative detection of SBV in 10 colonies belonging to three randomly selected apiaries showed the significant difference of SBV DNA per honeybee before and after period of dsRNA treatment *(p* = 0.000042), SBV relative DNA per bee ranged from 3.20 × 10^2^ ± 1.92 × 10^1^ to 7.79 × 10^3^ ± 1.13 × 10^1^ copies (Fig 4). However, no symptoms of SBV infection were observed in all the 78 colonies (Fig 3).

**Fig 4.**
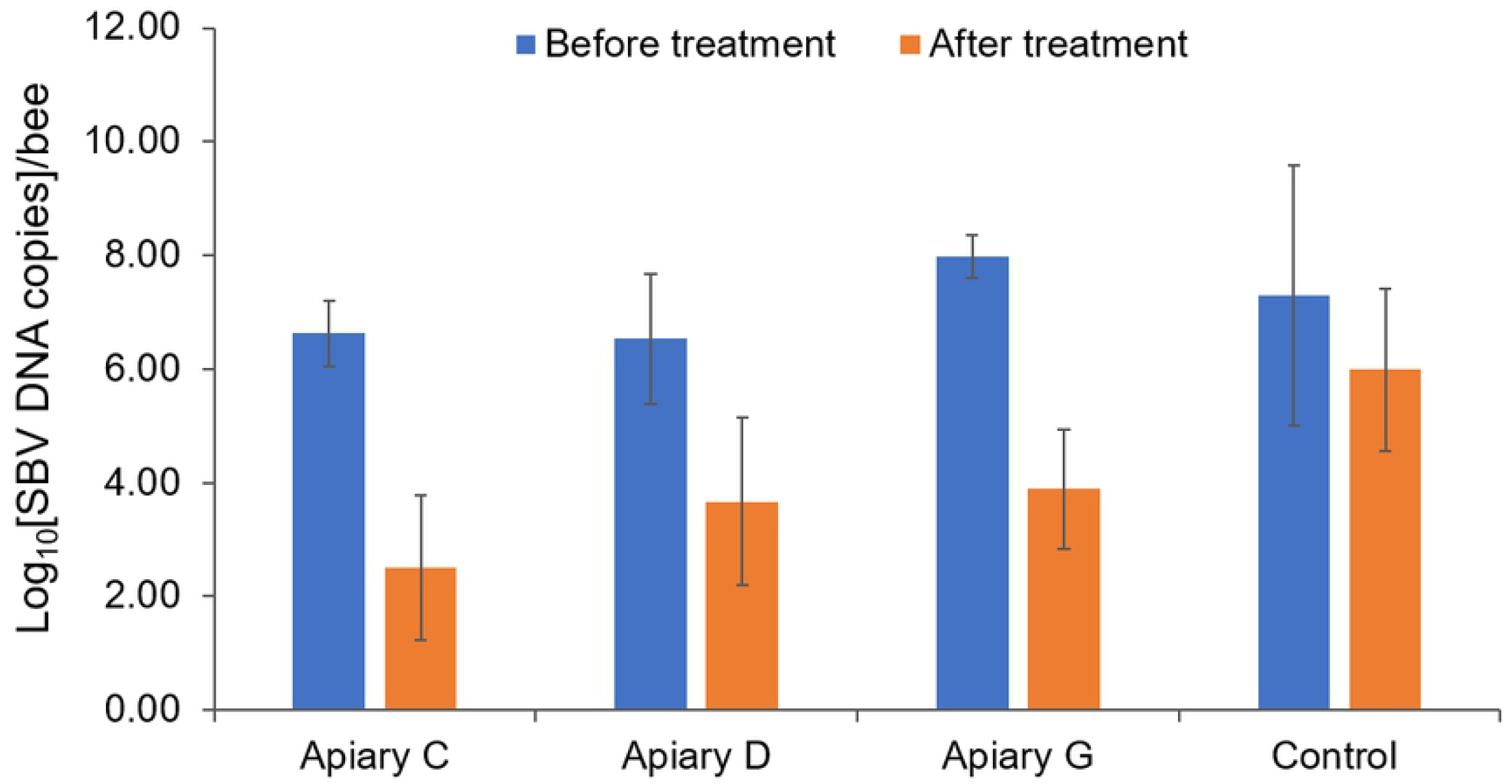
Quantitative detection of sacbrood virus (SBV) in double-stranded RNA (dsRNA) treatment apiaries. Ten colonies (from three apiaries, apiary C, D, and G) out of the 78 dsRNA treatment colonies belonging to 17 apiaries were randomly selected for quantitative detection of SBV before and after the 5^th^ dsRNA administration. Significant differences in SBV DNA in the colonies were observed (*n* = 10, *p* = 0.000042) before and after treatment. Meanwhile, the control colonies (*n* = 7) showed no significant differences (*p* = 0.077).

As for large-scale dsRNA application, the colonies with positive SBV detections by qRT-PCR but no clinical signs of disease were used as the prevention group, with application intervals of 2 or 4 weeks (Table 4). All the colonies exhibited healthy conditions during the period of inspection. However, in the group in which dsRNA was administered to colonies with SBV disease for treatment and without confining the queens to stop spawning during the treatment period, dsRNA was not effective or was only partially effective against SBV, depending on the application interval. dsRNA was administered to these colonies every 4 weeks, 2 weeks, or 1 week, which can be maintained for 2 months, longer than 2 months, or 8 months before the colony collapse occurred (Table 4).

**Table 4.**
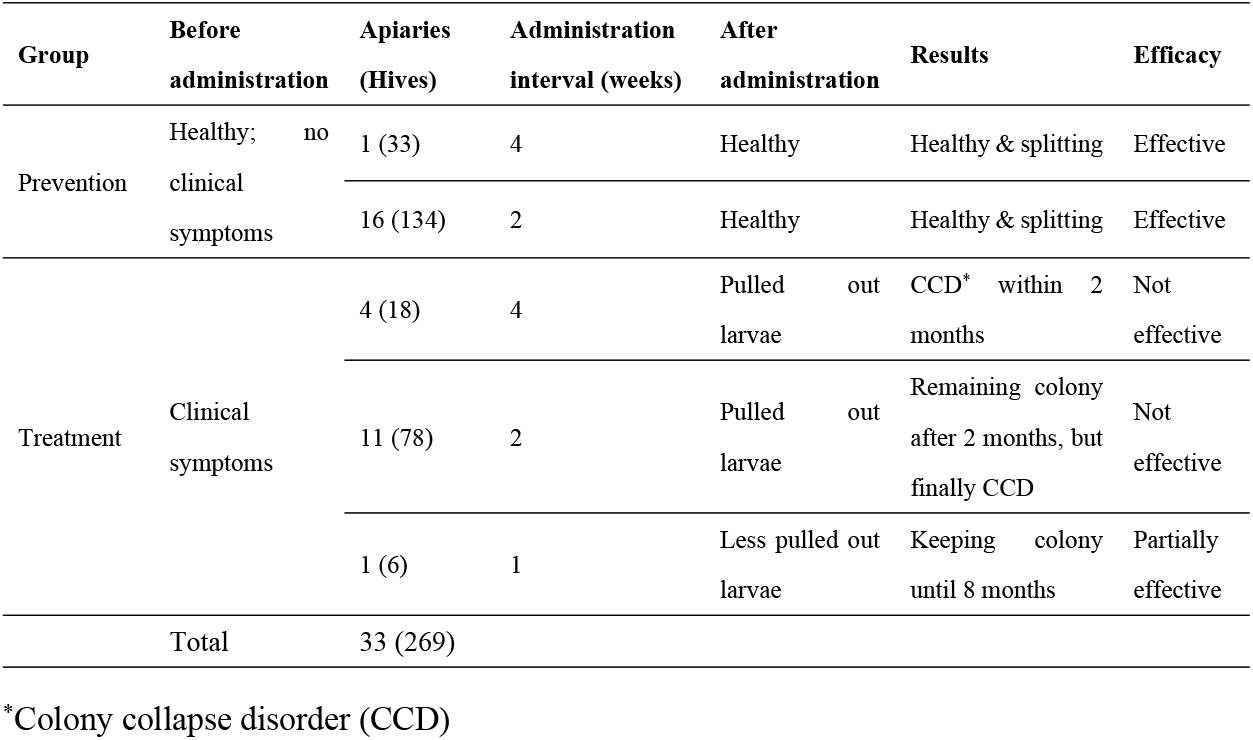
Extended field application of double-stranded RNA (dsRNA) for the prevention and treatment of sacbrood virus (SBV)

## Discussion

dsRNA for use against SBV was produced and tested at three levels: *in vitro* reared larvae, under controlled conditions, and for large-scale field application. The capacity of dsRNA to inhibit SBV was demonstrated as it was efficiently used as a gene therapy for the prevention and treatment of SBV. In addition, dsRNA was demonstrated to be safe for application in honeybees.

Viral diseases significantly increase the loss of honeybee colonies worldwide, with more than 20 viruses that infect honeybees being identified [27]. However, to our knowledge, there are no specific chemotherapies for the prevention and treatment of these viral diseases. Traditional methods for the management of viral diseases are good practice for beekeeping and help with the early diagnosis of diseases and selection of honeybee strains that are resistant to viruses [28]. When RNAi was demonstrated to effectively inhibit Israel acute paralysis virus (IAPV) and SBV [7, 23], a gene therapy relying on dsRNA for field treatment of IAPV was developed in 2010 [22]. However, the high costs of the *in vitro* production of dsRNA leads to difficulties in large-scale production. A method for the *in vivo* production of dsRNA against SBV was introduced in 2016 [24]. Although the cost was lower compared with that of *in vitro* production (224 USD per mg dsRNA), it was still highly expensive for field application, which requires around 20 mg for one dose. This cost-related restriction was addressed in the present study. The large-scale production of dsRNA in coordination with Genolution, Inc. (Seoul, South Korea) decreased commercial costs to around 10 USD for 60 mg, which was enough for three doses of field application.

Although SBV causes the death of infected larvae [2], the clinical signs in infected adult honeybees are unclear [29, 30]. Therefore, during the field application of dsRNA for the disease prevention in infected colonies, the queens were confined for the first 3 weeks so that the number of larvae in the colony did not increase any further. This mitigated the replication of SBV and would be helpful for dsRNA treatment in the infected colonies. After five administrations of dsRNA, the presence of SBV decreased below the level required to generate the disease in the infected colonies (≤7.79 × 10^3^ ± 1.13 × 10^1^ copies/bee) [31]. However, these results were obtained after 2 months of treatment, and further investigation to assess how long the administration of dsRNA could maintain SBV at a safe level, as well as the time required for SBV to reach the DNA replication level that causes the associated disease after terminating the use of dsRNA, is required.

The dsRNA present in the cells of honeybee could be processed into small interfering RNAs (siRNA). Such siRNA are then incorporated with the RNA induced silencing complex (RISC) [32]. The siRNA in RISC becomes a guided strand via complementary binding to target RNA and the RISC complex with Argonaute protein, a major component, causing the degradation of targeted RNA [33, 34]. Therefore, supplying the anti-SBV dsRNA to the honeybee prior to SBV exposure could provide protection from SBV infection. The effectiveness of dsRNA in preventing SBV disease was demonstrated following the administration of single dose of 10 mg 3 days before artificial infection with SBV, which prevented disease outbreak. The detection of SBV indicated that the virus remained at a high level 7 days following infection before decreasing considerably from days 7 to 10 and being completely eliminated after 3 weeks (data not shown). Furthermore, the five dsRNA administrations (20 mg/dose) at 1-, 2-, and 4-week intervals indicated 100% efficacy in protecting colonies from SBV disease outbreaks. However, the present study did not assess how long dsRNA can remain in honeybees to maintain protection following each administration. Understanding how long the dsRNA could confer protection could facilitate its large-scale application.

## Conclusions

The low production cost of dsRNA in the present study enabled the use of dsRNA as a specific drug for the prevention and treatment of SBV for beekeeping. The application of dsRNA was safe for honeybees, and the usefulness of dsRNA for preventing SBV outbreaks as well as for the treatment of SBV infection was demonstrated. However, further investigation is important to understand the duration of protection after each administration, the adequate administration frequency, and the optimal season for application.

## Acknowledgments

We would like to express our gratitude to Dae Rib Kim for his technical assistance for handling hives and developing dsRNA injection route.

## Supporting information

**S1 Fig. Detection and isolation of sacbrood virus (SBV) from infected honeybee larvae.** (**A**) SBV in infected *Apis cerana* larvae was detected with qRT-PCR and confirmed by electrophoresis, with an expected band of 131 bp in length. SBV+ and SBV-are samples with positive and negative detection, respectively. “+” and “-” are the positive and negative controls with SBV recombinant DNA and without a DNA template, respectively. (**B**) Band of purified SBV from infected larvae in 40% sucrose gradient density (indicated by the arrow). (**C**) Viral particles of SBV, with a size of 27.8 ± 0.4 nm and a shape similar to that of other picornaviruses, observed and measured under transmission electron microscopy.

**S2 Fig. Double-stranded RNA (dsRNA) of sacbrood virus VP1.** The dsRNA (185 ng) from one of the tubes containing dsRNA (685 mg/tube) was loaded into 1% agarose gel for electrophoresis. The 596 bp band corresponds to dsRNA. “M” is 1 kb DNA marker.

**S3 Fig. Standard curves of sacbrood virus (SBV) DNA amplification.** (**A**) Ten-fold serial dilution of SBV recombinant DNA, namely, from 10^10^ to 10^1^ copies, was used for triplicate PCRs. (**B**) Correlation between the initial number of DNA copies (log10-transformed values) and the cycle threshold of amplification.

## Notes

### Competing Interest Statement

The authors have declared no competing interest.

